## Supplementary Information for "Automated genome mining predicts structural diversity and taxonomic distribution of peptide metallophores across bacteria"

### Supplemental Discussion and Figures

Tables S1-S3 are located in the separate supplemental Excel (.xlsx) file

#### Supplemental discussion 1: Details of the detection strategy

Generally, draft pHMMs were built from alignments of known and predicted NRP metallophore biosynthesis genes collected from literature (Fig. S1). Initial bitscore cutoffs were determined by scanning MIBiG 2.0[14] gene clusters, as well as 38 additional experimentally characterized NRP metallophore BGCs that we added to MIBiG 3.0 (Supplemental Table 1).[15] Final cutoffs were determined by scanning 28,688 NRPS cluster regions from the antiSMASH database and manually inspecting hits near the initial cutoff to determine if they are likely true NRP metallophore BGCs based on the presence of genes encoding membrane transport, metal acquisition (ex: ferric reductases[16–18]), and multiple chelating groups. The pHMMs were iteratively refined where required by adding additional low-scoring putative true positives to the seed alignments until clear bitscore separations appeared between (putative) true and false hits (Fig. S1).

##### *Detection of catechols and phenols*

Like the aromatic amino acids, 2,3-dihydroxybenzoic acid (2,3-DHB) and salicylic acid are derived from chorismate (**Figure 1**B).[19–21] The three-step synthesis of 2,3-DHB is catalyzed by an isochorismate synthase (`EntC`), an isochorismatase (EntB), and a 2,3-dihydro-2,3-dihydroxybenzoate dehydrogenase (`EntA`).[19] The sole presence of `EntA` matches in a cluster was sufficient to detect 2,3-DHB-containing metallophore BGCs; however, only `EntC` could be used to eliminate the nearly-identical 3-hydroxyanthranilic acid pathway,[22] and thus both `EntA` and `EntC` were required to accurately detect 2,3-DHB production. Salicylate biosynthesis was detected by the presence of either an isochorismate pyruvate-lyase (`IPL`)[20] or a bifunctional salicylate synthase (`SalSyn`).[21] Two condensation domain subtypes specific to catecholic and phenolic metallophores were also included as independent detection rules: VibH-like enzymes (`VibH`), which condense 2,3-DHB to diamines and polyamines,[23,24] and tandem heterocyclization domains (`Cy_tandem`) are sometimes responsible for Ser, Thr, or Cys cyclization.[25]

##### *Detection of β-hydroxyamino acids*

An analysis of siderophores containing β-hydroxy-aspartate (β-OHAsp) and β-hydroxy-histidine (β-OHHis) delineated three families of siderophore-specific Fe(II)/α-ketoglutarate-dependent enzymes responsible for β-hydroxylation of Asp (`TBH_Asp` and `IBH_Asp`) or His (`IBH_His`).[26] More recently, β-OHAsp-containing siderophores named cyanochelins were isolated from a diverse group of cyanobacteria.[27] The BGCs contain two additional classes of β-hydroxylases, tentatively named **CyanoBH_Asp1** and **CyanoBH_Asp2**. Because of the polyphyletic nature of these hydroxylases, well-performing pHMMs could not be made using the standard workflow used for other pathways (**Figure S1**); even after many iterations, non-metallophore NRPs were captured as false positives, such as the β-OHAsp-containing lipopeptide turnercyclamycin.[28] Instead, pHMM construction was guided by a phylogenetic tree of β-hydroxylases from NRPS BGCs (**Figure S2**). Clades likely to be involved in metallophore biosyntheses were located using known genes from reported BGCs,[26,27] and by the presence of nearby metallophore-specific transporter families (*vide infra*). Several unreported clades of amino acid β-hydroxylases were identified that may also be involved in metallophore biosynthesis based on their co-occurrence with transporter genes (**Figure S2**). However, these putative β-hydroxylases were not included in the current NRP metallophore rules due to a lack of experimental evidence for metallophore production. A negative constraint (in the form of a competing pHMM for **SBH_Asp**) was used to exclude the syringomycin family of BGCs, which contain a clade of β-hydroxylases that sit within the IBH_Asp clade.[26]

##### *Detection of hydroxamic acids*

Hydroxamates are all produced by the hydroxylation and acylation of a primary amine. In peptidic metallophores, the amine is usually the sidechain of ornithine (Orn) or Lys, which is first hydroxylated by a flavin-dependent monooxygenase (**Orn_monoox** or **Lys_monoox**, respectively). Vicibactin is an unusual hydroxamate siderophore[29] that could not be accurately captured by either **Orn_monoox** or **Lys_monoox**. The Orn hydroxylase VbsO is more similar to those found in NIS pathways, and a VbsO pHMM was non-specific. Instead, the acyl-hydroxyornithine epimerase[29] **VbsL** is used to detect vicibactin. In two non-metallophore pathways,[30,31] hydroxyornithine or hydroxylysine are the substrates for N-N bond-forming enzymes. No clear bitscore separation could be achieved to eliminate these false positives, so two negative constraints were added: the presence of **KtzT** associated with biosynthesis of piperazates,[30] and **MetRS-like**, associated with several hydrazines.[31] Unfortunately, false positives may still arise if the ornithine monooxygenase and the negative constraint genes are (distantly) located within the same BGC region (>30 kbp), such as in the himastatin locus (MIBiG BGC0001117).

##### *Detection of other chelating groups*

Although the biosynthesis of the chelating amino acid graminine has not been fully elucidated, gene knockouts and stable isotope studies have revealed two enzymes, **GrbD** and **GrbE**, responsible for diazeniumdiolate formation from arginine.[32,33] The quinoline chelator Dmaq was first identified in anachelins,[34] and the biosynthesis was recently established for fabrubactins.[35] Synthesis is initialized by **FbnL** and **FbnM**, which form a two-protein heme peroxidase that oxidizes L-tyrosine to L-DOPA.[35] GrbDE and FbnLM were previously used as a handle for genome mining,[32,35] and our pHMMs gave similar results. We did not include detection of a pathway currently only reported in fabrubactins that produces two α-hydroxycarboxylate chelating moieties (Fig. 1A, bolded atoms).[31] Both substructures are proposed to be synthesized by the flavin-dependent monooxygenase FbnE. A profile HMM was built for FbnE (Supplemental dataset) but was not included in antiSMASH, as all BGCs with **FbnE** hits were either captured independently by **FbnL** and **FbnM** (Dmaq biosynthesis), or appeared to be false positives based on a lack of other metallophore-related genes. The diverse pyoverdine family of siderophores is defined by a fluorescent chelating chromophore (**Figure 1**). Chromophore maturation is driven by the tyrosinase **PvdP** and the oxidoreductase **PvdO**.[36,37] The two genes are sometimes in separate loci, and including both as independent pHMMs increased the number of pyoverdine BGCs detected. These pHMMs also capture the biosynthesis pathway for azotobactin, which contains a slightly different chromophore (**Figure 3**).[38]

**Figure S1. Workflow for developing an NRP-metallophore-specific profile hidden Markov model (pHMM) and significance score cutoff for an enzyme (sub-)family.** (1) Examples are collected from literature, the amino acid sequences are aligned with MUSCLE and (2) a pHMM is constructed with HMMER3. (3) BGCs of known function from MIBiG are scanned for matches to the pHMM to generate a preliminary bitscore cutoff. (4) NRPS BGC regions from the antiSMASH database are scanned for matches to the pHMM and sorted by bitscore. (5) Starting at the bitscore cutoff, BGCs are manually annotated to predict if they encode the biosynthesis of NRP metallophores using features such as genes encoding membrane transport, metal acquisition (ex: ferric reductases), and the biosynthesis of multiple chelating groups. In a properly functioning system, the bitscore cutoff delineates putative true and false positives to accurately detect the NRP-metallophore-related enzymes (bottom right). However, a bitscore cutoff adjustment may be required (bottom middle), or low-scoring true positives may need to be added to the pHMM seed alignment in an iterative process (bottom left).

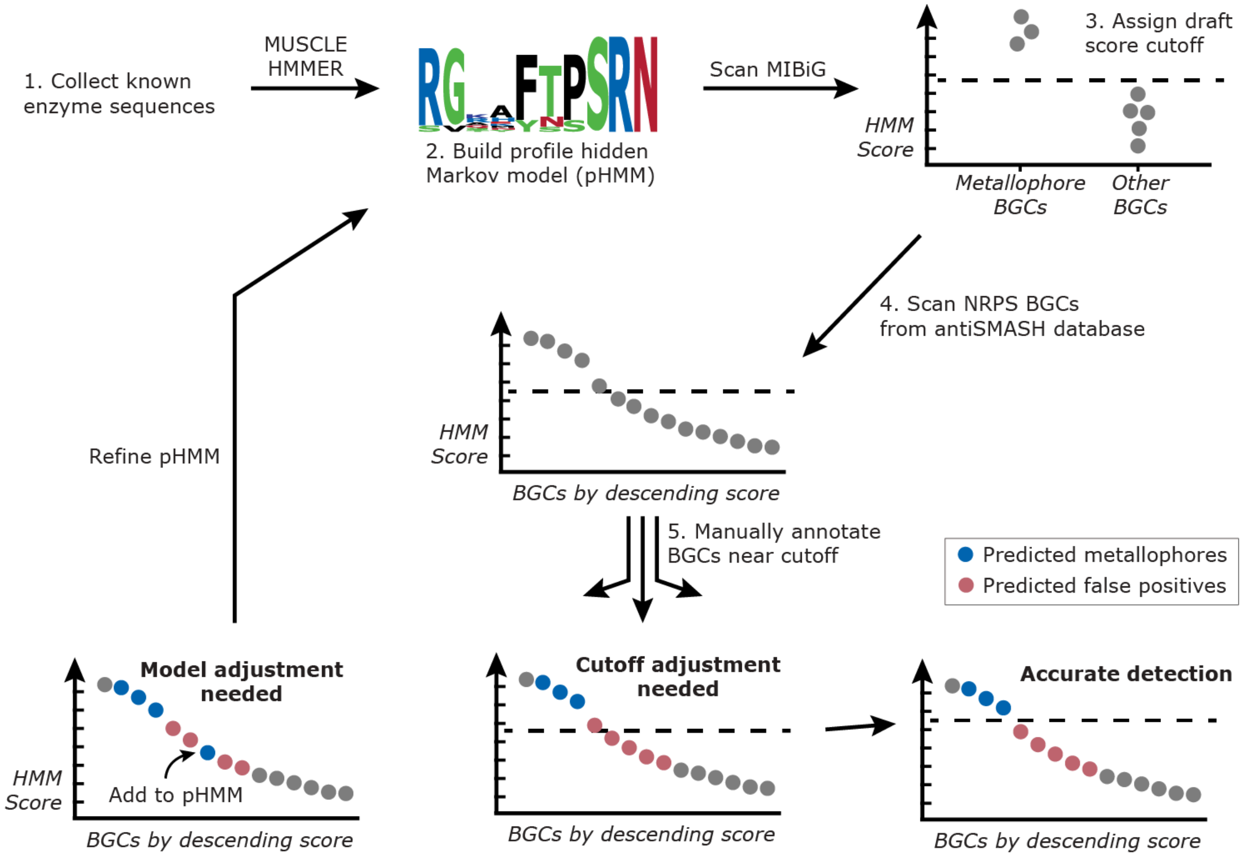

**Figure S2. A maximum-likelihood phylogeny of putative β-hydroxylases found in NRPS BGC regions.** β-Hydroxylase subtypes found in characterized siderophore BGCs are highlighted and labeled; the red SBH_Asp clade consists of non-metallophore phytotoxins.[1] Gray clades indicate possible metallophore β-hydroxylases that currently have no experimentally characterized representative. Amino acid sequences similar to known siderophore β-hydroxylase subtypes were extracted from the antiSMASH database (v3) and dereplicated prior to tree reconstruction. The right-hand bar gives the number of unique siderophore-related transporter families[2] present in the BGC region containing the β-hydroxylase.

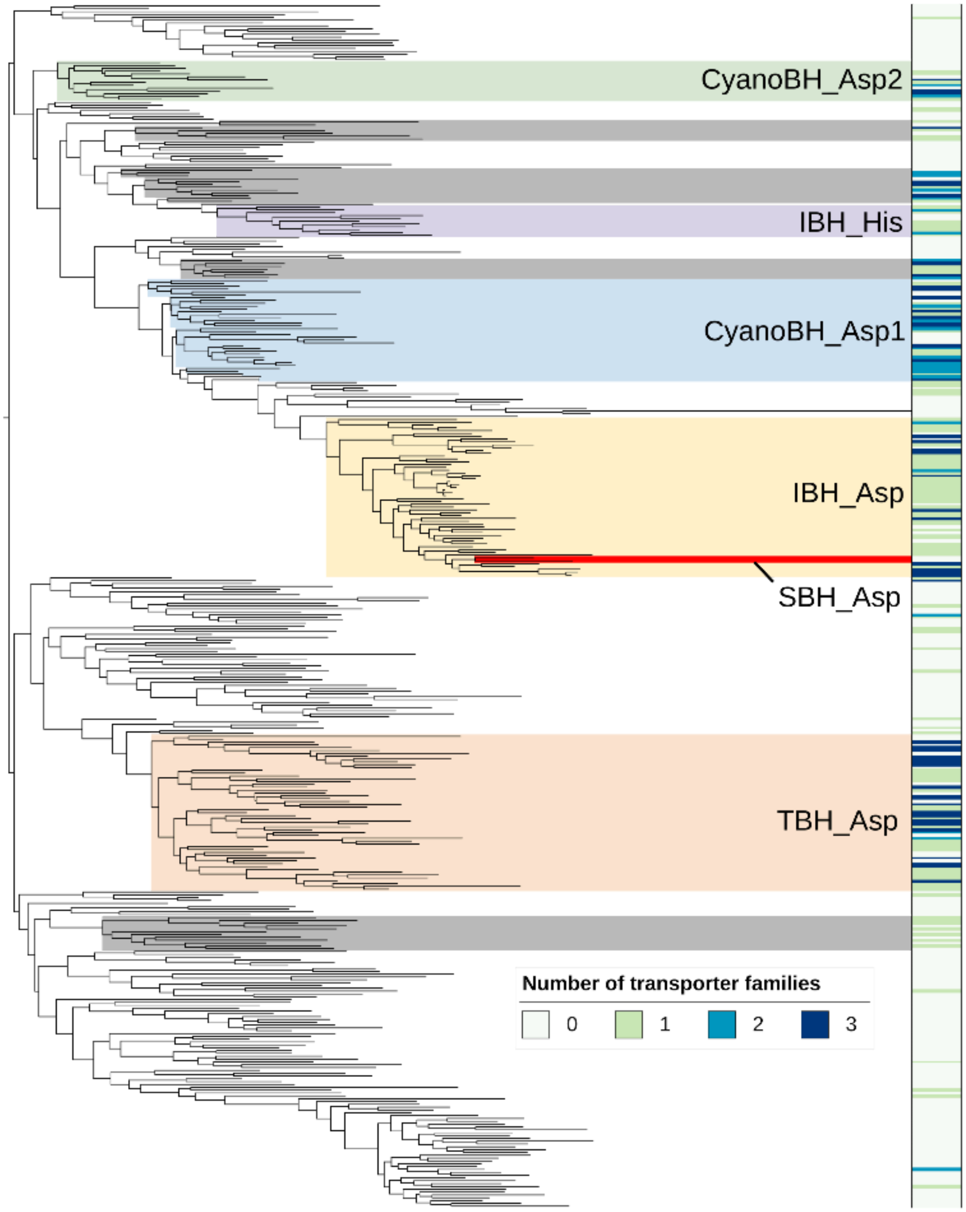

#### Supplemental discussion 2: A BGC from *Sporomusa termitida* putatively encoding for a menaquinone-related chelating group

One particularly promising novel BGC was found in the genome of *Sporomusa termitida* DSM 4440, an anaerobic acetogen (Figure S3). The BGC was manually annotated as a putative metallophore due to the presence of a TonB-dependent outer membrane receptor, as well as NRPS domains, methyltransferases, and oxidoreductases homologous to those involved in the biosynthesis of pyochelin and other thiazol(id)ine metallophores. A salicylate synthase gene, present in similar clusters, appears to have been replaced by a partial menaquinone pathway (*menFDEB*). We thus predicted that the salicylate moiety would be replaced by 1,4-dihydroxy-2-naphthoic acid (Figure S3). The Natural Product Atlas contained one family of bacterial compounds with that substructure, karamomycins, which were isolated from an unsequenced strain of *Nonomuraea endophytica* (Figure S3).[42] Karamomycins also contain thiazol(id)ines, as predicted for the *S. termitida* DSM 4440 BGC product. Hence, karamomycins are likely produced by a homologous BGC, and both compounds may be involved in trace metal binding and transport.

**Figure S3 (next page). Analysis of a novel putative NRP metallophore BGC from *Sporomusa termitida* DSM 4440.** (A) Clinker comparison of the *S. termitida* BGC with homologous loci. The *Sporomusa* sp. KB1 BGC contains a salicylate synthase gene detectable by the new antiSMASH rules, while *S. termitida* DSM 4440 instead contains several genes homologous to the menaquinone locus of *Desulfitobacterium* spp.[3] The cluster comparison was generated with clinker v0.0.26.[4] (B) A proposed metallophore biosynthesis pathway encoded by the *S. termitida* DSM 4440 BGC. 1,4-Dihydroxy-2-naphthoic acid is synthesized from chorismic acid by homologs of MenFDHBE, encoded by SPTER_RS05985-06005, and MenC, encoded elsewhere in the genome (SPTER_RS21050). The C4 phenol is likely methylated by O-methyltransferase SPTER_RS06015. The naphthoic acid moiety of Karamomycin C (inset box), characterized from an unsequenced strain of *Nonomuraea endophytica*,[5] is predicted to be synthesized by a similar pathway. Five NRPS genes in *S. termitida* DSM 4440 encode for the biosynthesis of the core structure by the condensation and cyclization of four Cys residues. The C-methyltransferase of SPTER_RS06025 is predicted to be inactive, as observed in the homologous ulbactin pathway.[6] The completed scaffold is released by thioesterase SPTER_RS05970, possibly producing the same tricyclic substructure observed in karamomycin C and ulbactin F (inset box). The final predicted structure accounts for the actions of two thiazoline reductases (SPTER_RS05950 and SPTER_RS05980) and a methyltransferase (SPTER_RS05955), although the regiochemistry and timing of these transformations is unclear.

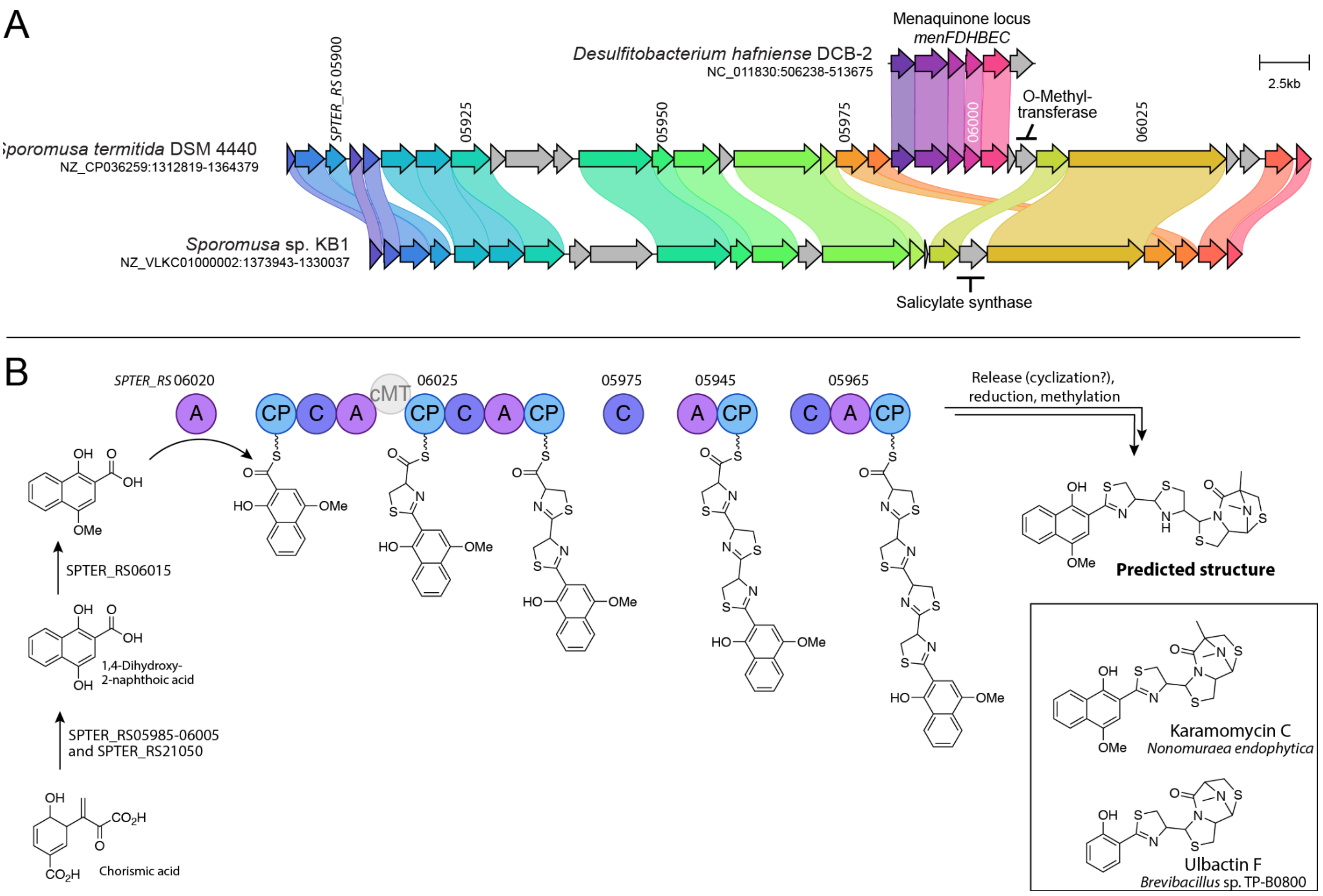
A
Sporomusa termitida DSM 4440
NZ_CP036259:1312819-1364379
Sporomusa sp. KB1
NZ_VLKC01000002:1373943-1330037
Desulfitobacterium hafniense DCB-2
NC_011830:506238-513675
Menaquinone locus
menFDHBEC
SPTER_RS 05900
05925
05950
05975
06000
06025
O-Methyl-
transferase
Salicylate synthase
2.5kb
B
SPTER_RS 06020
06025
05975
05945
05965
cMT
Release (cyclization?),
reduction, methylation
Predicted structure
SPTER_RS06015
1,4-Dihydroxy-
2-naphthoic acid
SPTER_RS05985-06005
and SPTER_RS21050
Chorismic acid
Karamomycin C
Nonomuraea endophytica
Ulbactin F
Brevibacillus sp. TP-B0800

**Figure S4**. **Similarity network of complete NRP metallophore BGC regions from RefSeq representative genomes.** Nodes are colored blue if they belong to a GCF with a reference BGC, and orange if they are dissimilar from any reference BGC. The BiG-SCAPE network is identical to that in Figure 3 and reference BGC numbering corresponds to Supplemental Table

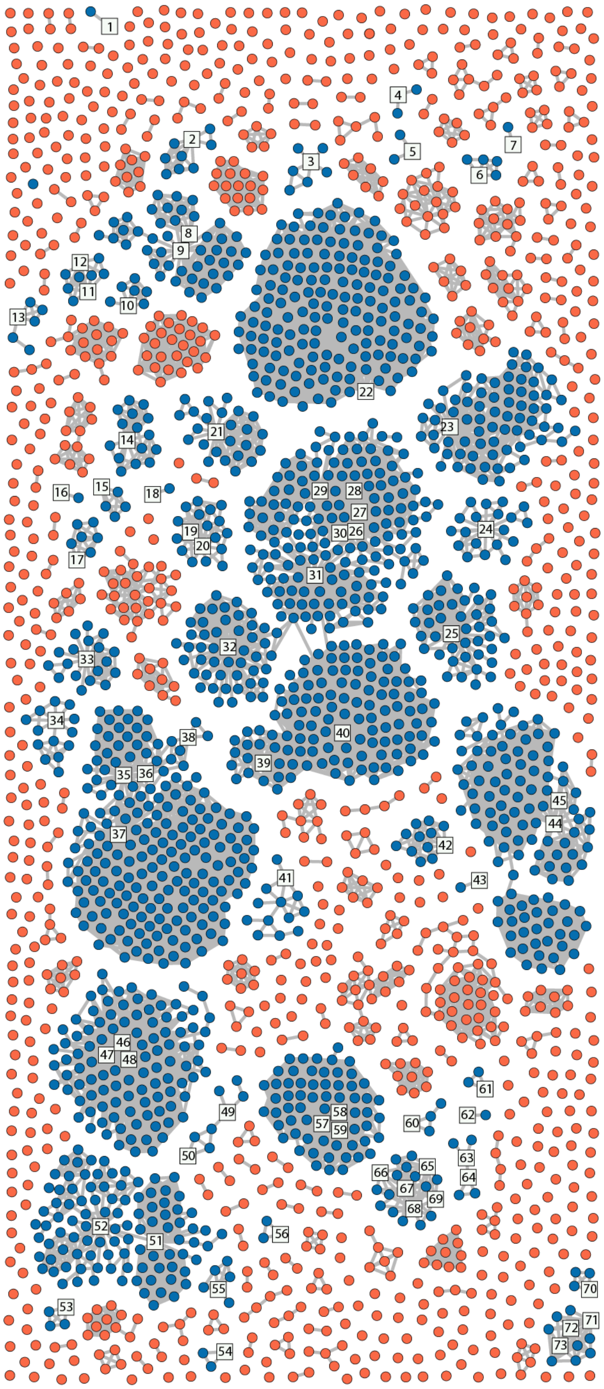

1.

#### Supplemental methods and results for siderophore isolations

##### Methods

All strains were obtained from the Leibniz Institute DSMZ. Unless otherwise stated, all water is doubly deionized (dd $H_2O$), 18 MΩ.

*Terasakiispira papahanaumokuakeensis* DSM 29361 and *Buttiauxella brennerae* DSM 9396 were cultivated in a 4L Erlenmeyer flask acid washed with 4M HCl. *T. papahanaumokuakeensis* was grown in 2L of the following medium: glycerol phosphate disodium 4 g/L, NaCl 25 g/L, KCl 0.67 g/L, $CaCl_2$ * 2 $H_2O$ 1.36 g/L, $NH_4Cl$ 4 g/L, L sodium succinate hexahydrate 5 g/L and 25 mM MOPS pH 7.5. For *B. brennerae*, the medium was 2L: $K_2HPO_4$ 7 g/L, $KH_2PO_4$ 2 g/L, NaCl 0.6 g/L, $MgSO_4$ * 7 $H_2O$ 1 g/L, $NH_4SO_4$ 1 g/L sodium succinate hexahydrate 5 g/L. After autoclaving 20mL of filter sterilized D-glucose 50% (w/v) was added, completing the medium. The cultures were incubated at room temperature on an orbital shaker, 180 RPM. After 41h and 48h, the cultures of *T. papahanaumokuakeensis* and *B. brennerae*, respectively, were centrifuged at 6,000xg for 10 min at 4°C. The supernatant was decanted and approximately 100 g of XAD-4 resin added. The solution was agitated gently at room temperature for approximately 4 hours. Organic compounds were eluted from the resin with 100% methanol, which after rotary evaporation yielded the crude extract. Four catechol compounds produced by *B. brennerae* were isolated using reverse phase (RP) HPLC. The HPLC system was comprised of a YMC AQ12S05-2520WT column connected to two Waters 515 HPLC pumps and a waters 2487 dual wavelength absorbance detector. The mobile phase consisted of dd $H_2O$ and methanol, both amended with 0.05% trifluoroacetic acid (w/v). To separate the compounds a method of 10-100% methanol, 3% per min was utilized. The crude extract of *T. papahanaumokuakeensis* and the catechol compounds of *B. brennerae* were analyzed by ESI-MS and MS/MS on a Waters XevoG2-XS QTof instrument in positive mode electrospray ionization coupled to an ACQUITY UPLC H-Class system with a Waters BEH C18 column.

*Pseudomonas brassicacearum* DSM 13227 was cultured in a modified M9 minimal medium composed of 3.0 g/L disodium hydrogen phosphate heptahydrate, 1.5 g/L potassium dihydrogen phosphate, 2.5 g/L sodium chloride, 0.50 g/L ammonium chloride, 2.5 g/L disodium succinate, and 5.0 g sodium pyruvate in ultrapure (e.g., 18 mW) water, amended after autoclave sterilization with 20 mL/L 50% w/v Steri-filtered glucose solution, 1 mL/L 1M magnesium chloride, and 0.05 mL/L 1M calcium chloride. Culturing glassware was rinsed with 4M HCl before use to remove adsorbed iron(III). Seed cultures were inoculated in Luria-Bertani (LB) broth with single colonies of *P. brassicacearum* DSM 13227 grown on LB agar and grown for at least 24 h at 30°C. Cultures were monitored by $OD_{600}$, and the culture supernatant was harvested at late log/early stationary phase ($OD_{600}$ ~ 1.2 au) with a positive Fe(III)-CAS response. Culture supernatants were obtained by centrifugation at 6500 rpm for 20 min at 4 °C. To extract the siderophores, the culture supernatant was decanted and shaken with 100 g of Amberlite XAD-4 resin. The XAD-4 resin was prepared by washing with methanol and then equilibrating with ultrapure water. The resin and supernatant were orbitally shaken to equilibrate for 2 h at 150 rpm. The resin was filtered from the supernatant and washed with 0.5 L of ultrapure water. The adsorbed organics were eluted with 80% aqueous methanol. The eluent was concentrated under vacuum and stored at 4 °C. Ornicorrugatin and the pyoverdine siderophore were identified by UPLC-MS and LC-MS/MS analysis of the concentrated eluent.

**Results**

The crude supernatant extract from the predicted enterobactin producer *B. brennerae* was subjected to semi-preparative reverse-phase high-pressure liquid chromatography (HPLC) coupled to a UV/visible spectrophotometer. Four catecholic compounds were identified by their characteristic absorbance of UV light at 310 nm (Fig. S5). Their relative retention times were consistent with compounds found in supernatant extracts of the known enterobactin producer *Escherichia coli*, which produces the enterobactin fragments 2,3-DHB–Ser (DHBS), $(DHBS)_2$ and linear $(DHBS)_3$, in addition to enterobactin (cyclic $DHBS_3$; Fig. 4B).[47] The four compounds were purified and subjected to ultra-performance liquid chromatography (UPLC) coupled to electrospray ionization mass spectrometry (ESI-MS) (Figs. S6 and S7). The molecular ions of the purified compounds were each consistent with the enterobactin family (*m/z* 242.0647, 465.1157, 688.1614, and 670.1469, respectively).

Marinobactins A-E, differing in the identity of their fatty acid tails (Fig. 4A), were previously isolated from *Marinobacter* spp.[48] We searched for the compounds in the crude extract of *T. papahanaumokuakeensis* by UPLC-ESI-MS. Molecular ions consistent with all five of the marinobactins were detected (*m/z* 932.4986, 958.5159, 960.5315, 986.5422, and 988.5421, respectively, Figs 4C and S7). The crude extract was then analysed by tandem ESI-MS/MS, yielding fragmentation for the peaks putatively corresponding to marinobactins A-D (the peak at *m/z* 988.5421, putatively marinobactin E, was low abundance and did not give a clear spectrum). Each had similar fragmentation patterns, with $b_4$, $b_5$, $y_1$, and $y_2$ fragments further supporting the assignments of the marinobactins (Figs. S8-11). No peaks consistent with enantio-pyochelin (*m/z* 324.4; Fig. 1a) could be observed.

Siderophores were likewise identified from the crude extract of *P. brassicacearum* by UPLC-ESI-MS and -MS/MS. As hypothesised, ornicorrugatin (Fig. 4A) was indeed identified by searching the chromatogram for the expected molecular weight (*m/z* 1012.5). A candidate peak was identified with a singly, doubly, and triply charged molecular ion consistent with ornicorrugatin (*m/z* 1012.46, 506.74, and 338.17, respectively, Figs. 4D and S13). The tandem MS/MS spectrum closely matched that previously reported for ornicorrugatin (Fig. S14 and Table S4).[44] A chromatographic peak corresponding to pyoverdine was identified by the characteristic absorbance of the chromophore at 400 nm. A candidate compound was identified with singly and doubly charged molecular ions at *m/z* 1134.41 and 567.72, respectively (Figs. 4D S15). As predicted from the BGC analysis, this mass was identical to pyoverdine A214, the previously characterized siderophore of *P. brassicacearum* subsp. *brassicacearum* NFM 421 and *Pseudomonas* sp. A214 (Fig. 4A).[45,46] MS/MS fragmentation yielded B-fragments identical to those previously reported (Fig. S16).[45]

**Figure S5. Semi-preparative RP HPLC chromatogram of *B. brennerae* DSM 9396 crude extract.** The 4 peaks indicated by the arrows displayed catechol absorbance and were isolated

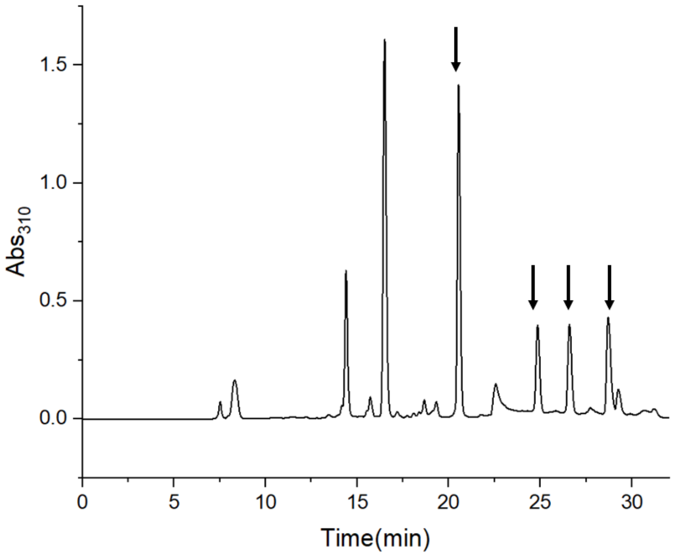

**Figure S6. UPLC-ESI-MS TIC of the purified catechol compounds.** From top to bottom, the molecular ions of the compounds were 670.1469 *m/z,* 688.1614 *m/z,* 465.1157 *m/z,* and 242.0647 *m/z*, which are consistent with enterobactin, the linear 2,3-DHB–Ser trimer, the 2,3-DHB–Ser dimer, and 2,3-DHB–Ser, respectively.

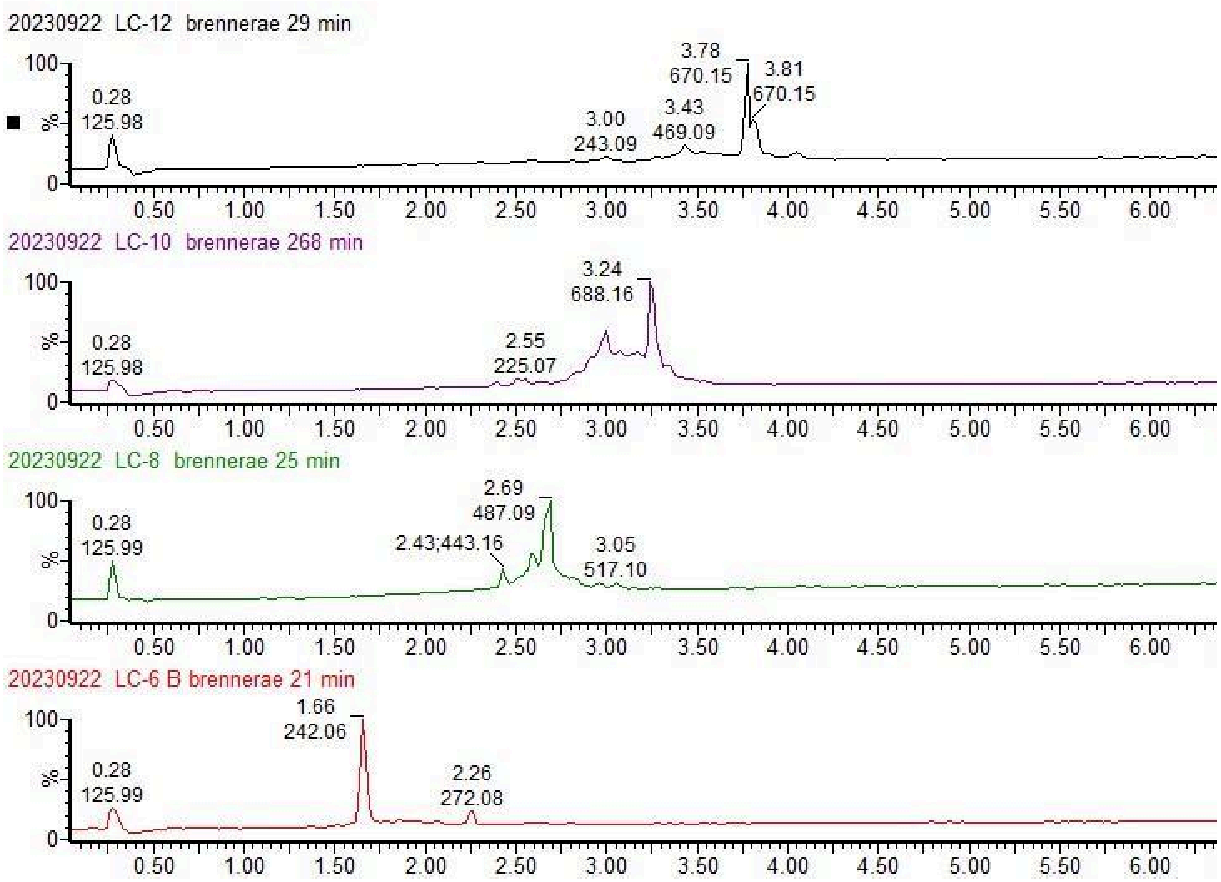

**Figure S7. Mass spectra of the purified catechol compounds.** From top to bottom, the molecular ions of the compounds were 242.0647 *m/z*, 465.1157 *m/z,* 688.1614 *m/z* and 670.1469 *m/z* which is consistent with 2,3-DHB–Ser, the 2,3-DHB–Ser dimer, the 2,3-DHB–Ser linear trimer, and enterobactin, respectively.

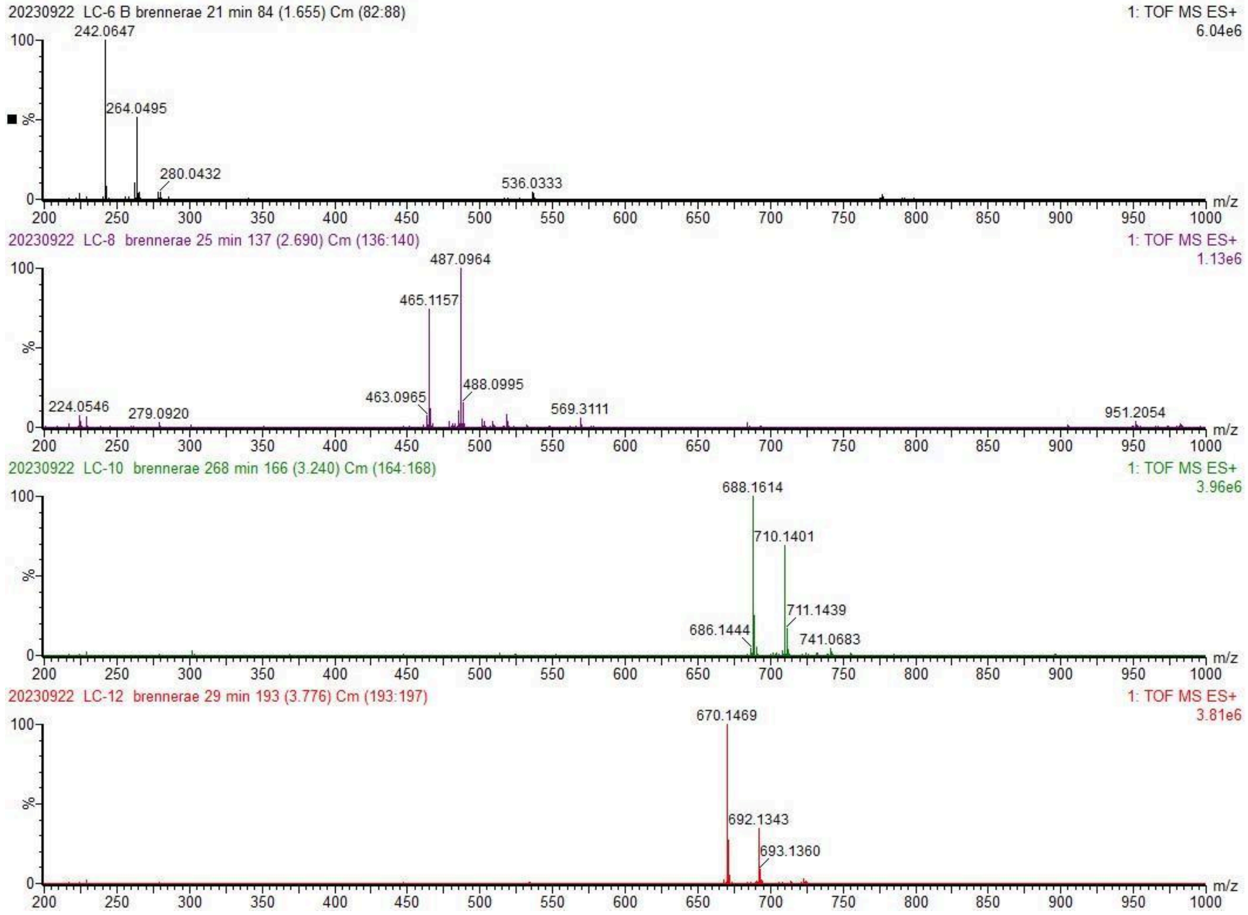

**Figure S8. UPLC-ESI-MS TIC the peaks with molecular ions consistent with marinobactin A-E are labeled (top).** Mass spectra of the labeled peaks in the TIC (bottom). The major peaks present are consistent with *m/z* value of Marinobactin A-E, 932.4986 *m/z*, 958.5095 *m/z*, 960.5315 *m/z*, 986.5422, and 988.5549 *m/z*, respectively.

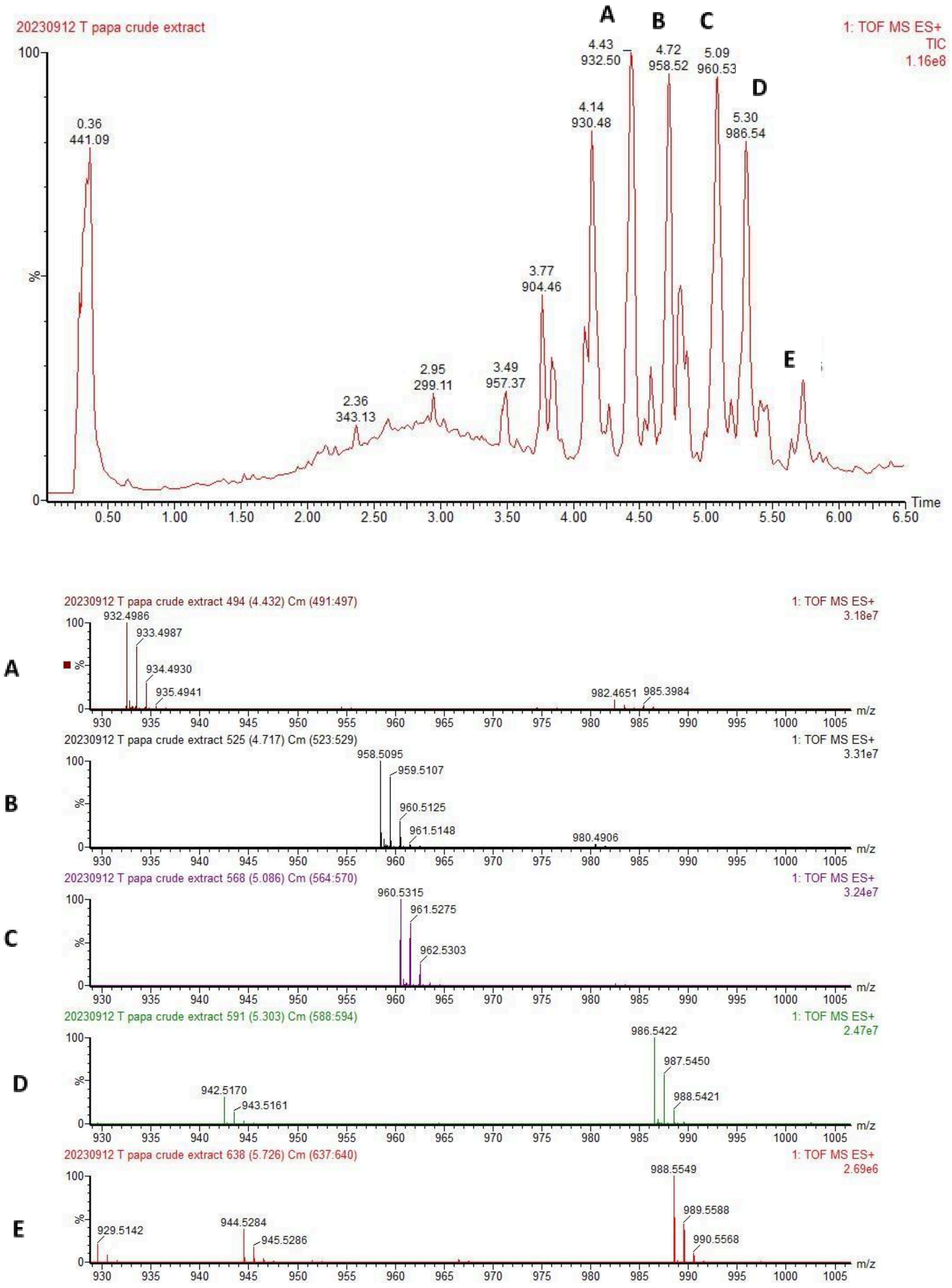

**Figure S9. ESI-MS/MS fragmentation of marinobactin A [M+H]+, 932 *m/z*.** The b fragments 655.3563 *m/z* and 742.3990 *m/z* support the assignment of marinobactin A.

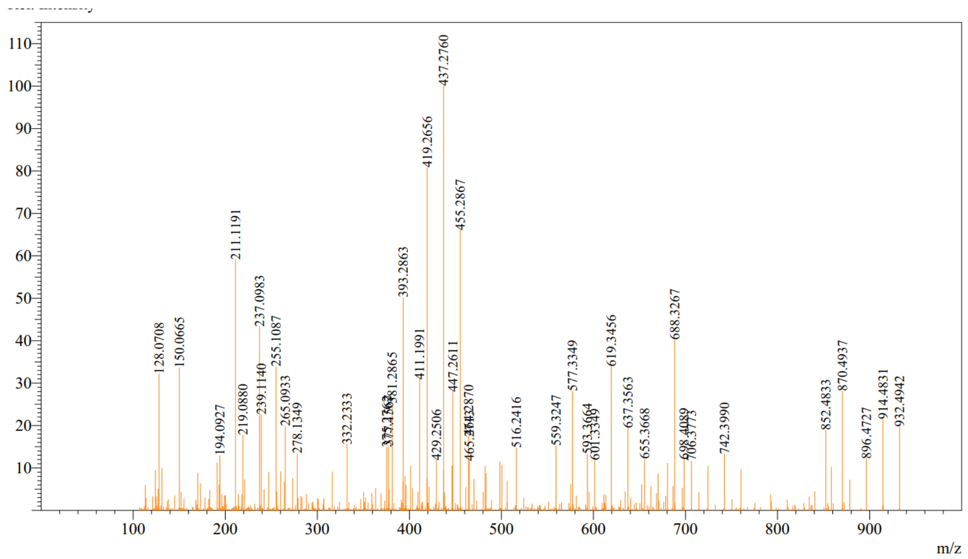

**Figure S10. ESI-MS/MS fragmentation of marinobactin B [M+H]+, 958 *m/z*.** The b fragments 681.3822 *m/z* and 768.4156 *m/z* support the assignment of marinobactin B.

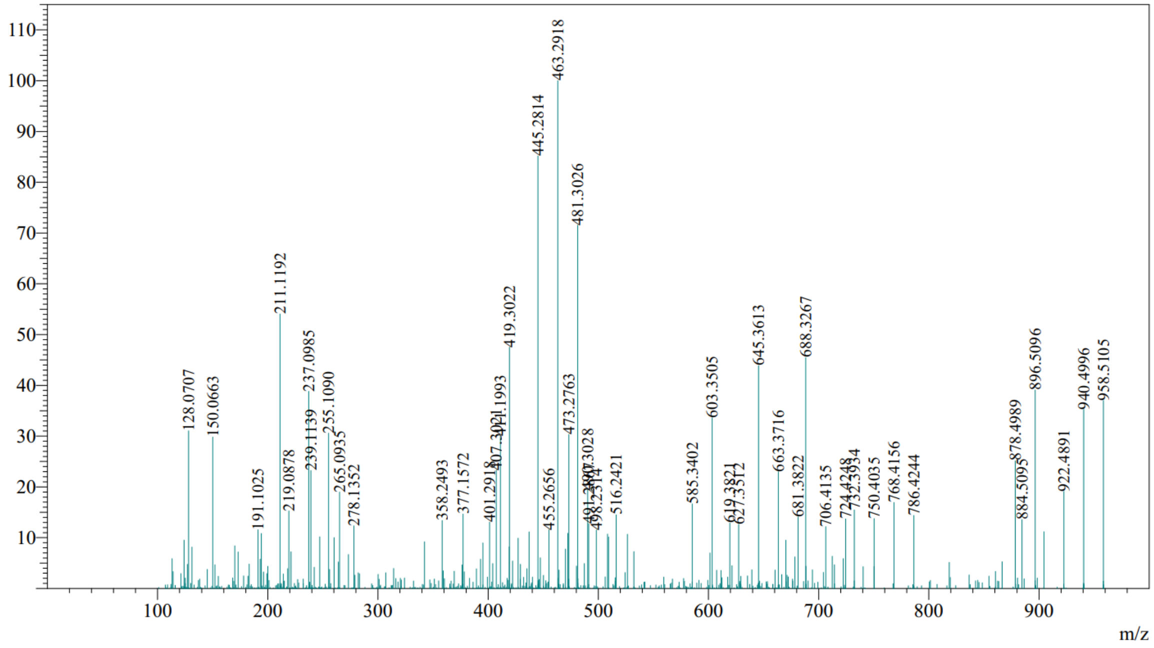

**Figure S11. ESI-MS/MS fragmentation of marinobactin C [M+H]$^+$, 960 *m/z*.** The b fragments 681.3985 *m/z* and 768.4298 *m/z* support the assignment of marinobactin C.

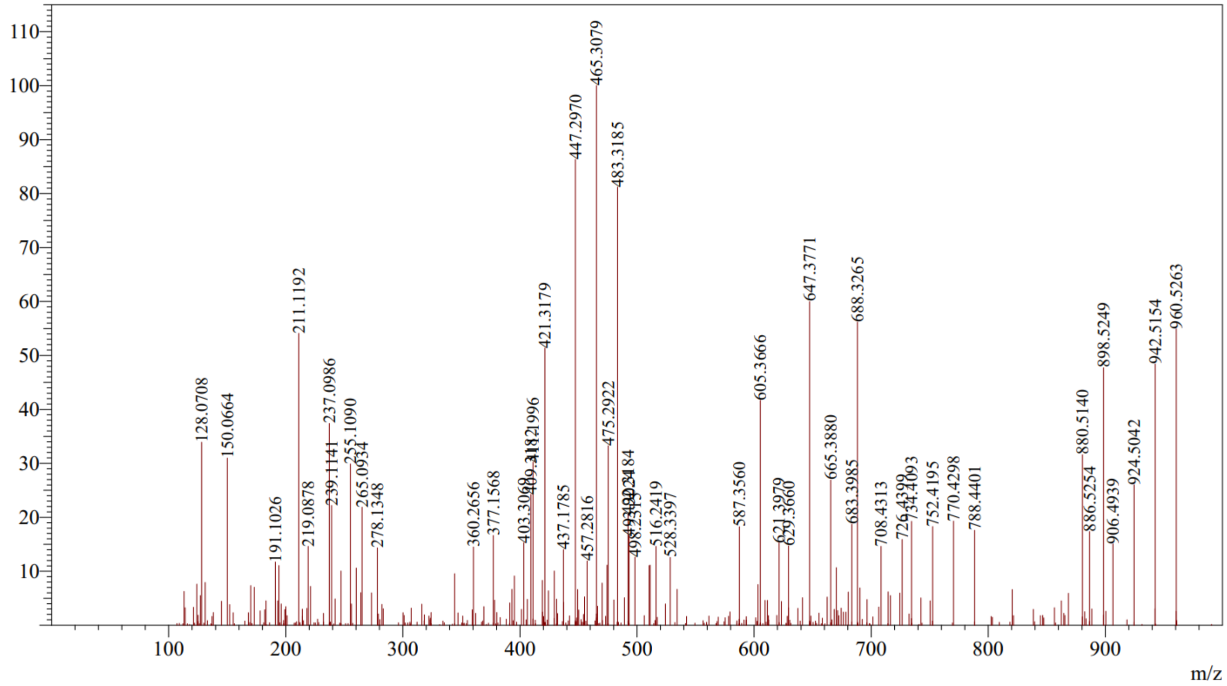

**Figure S12. ESI-MS/MS of marinobactin D [M+H]$^+$, 986 *m/z*.** The b fragments 709.4139 *m/z* and 796.4473 *m/z* support the assignment of marinobactin D.

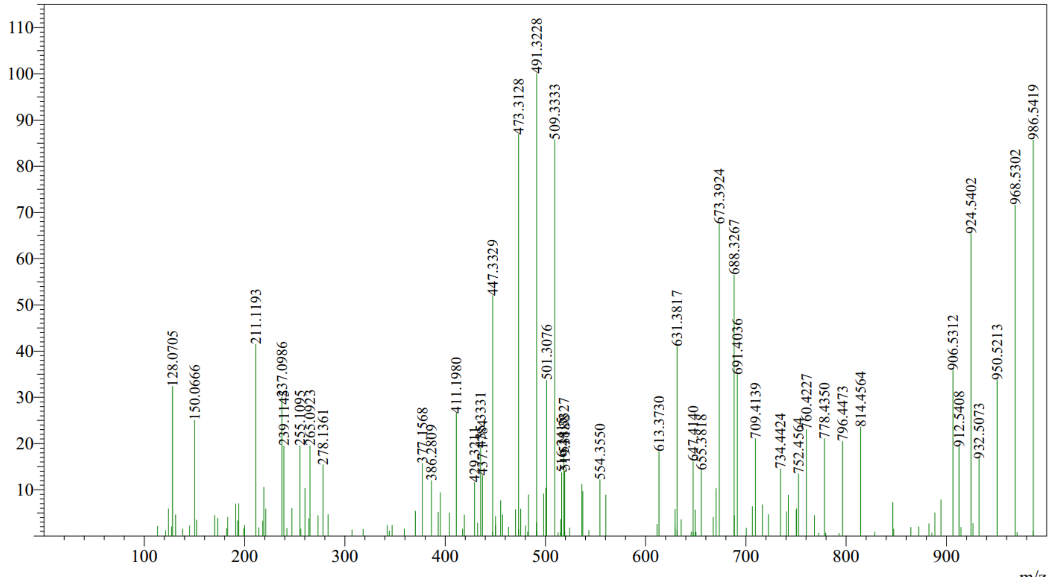

**Figure S13. UPLC-MS spectrum of ornicorrugatin siderophore from *Pseudomonas brassicacearum* DSM 13227.** The singly charged mass of *m/z* 1012.4 Da $[M+H]^{+}$ and the doubly charged mass, *m/z* 506.7 Da $[M+2H]^{2+}$, are consistent with the mass of the charged ornicorrugatin siderophore, *m/z* 1012.4699 Da $[M+H]^{+}$.

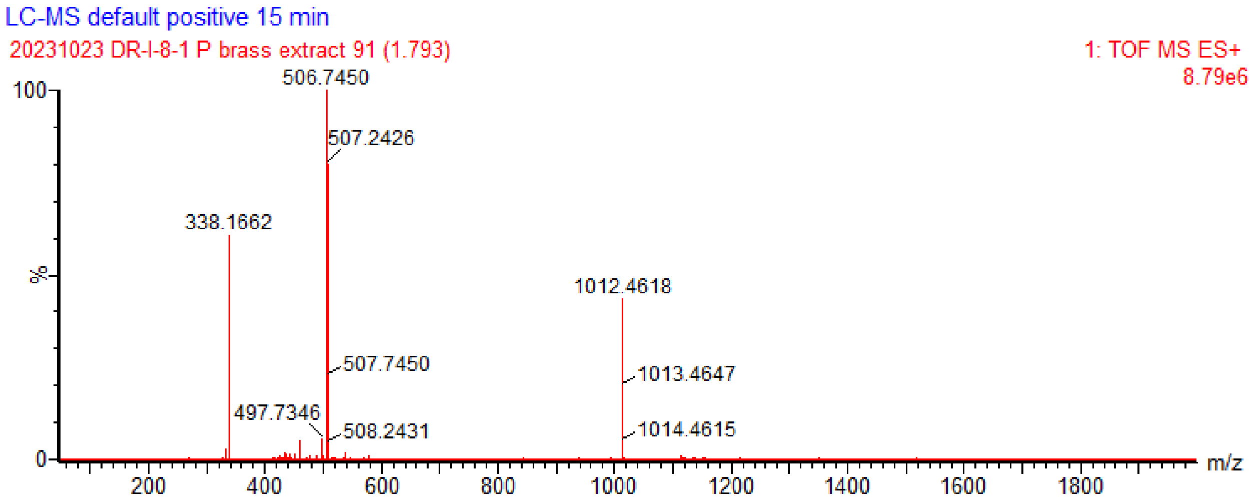

**Figure S14. LC-MS tandem MS spectrum of ornicorrugatin siderophore from *Pseudomonas brassicacearum* DSM 13227.** Average collision energies of (top) 60 eV and (bottom) 35 eV.

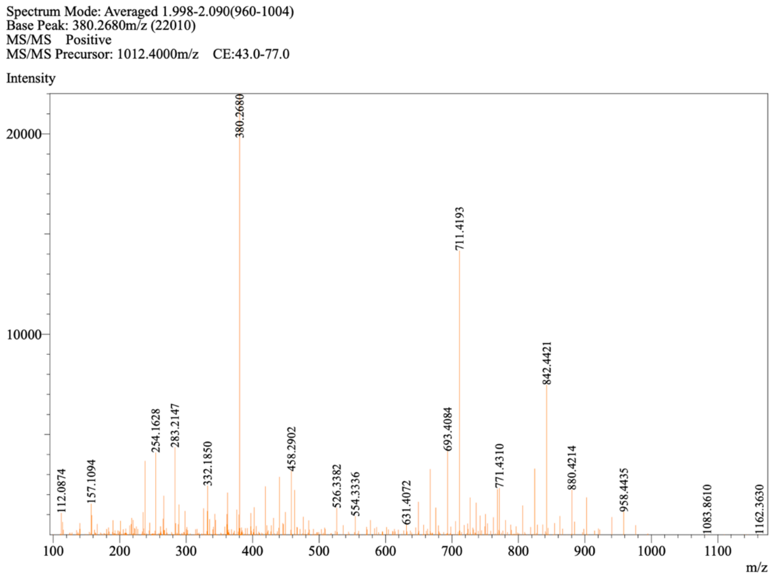

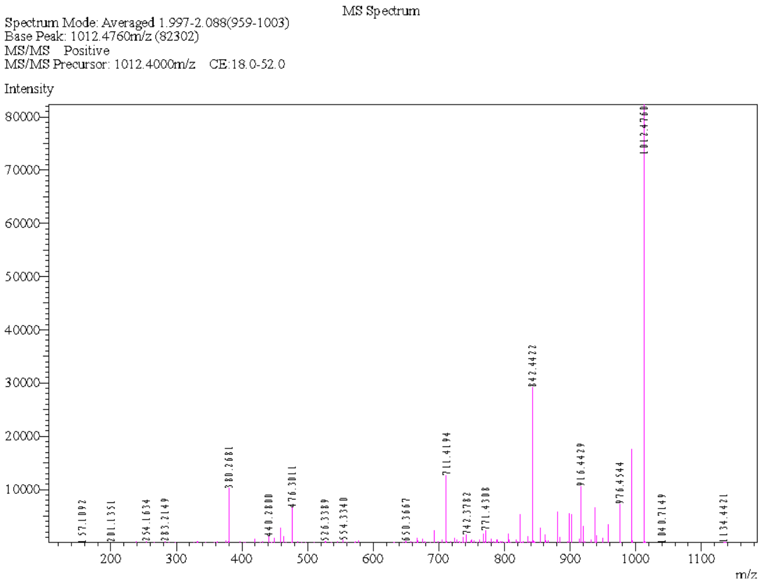

**Table S4: Selected fragment ions observed for ornicorrugatin produced by *Pseudomonas brassicacearum* DSM 13227.** All given fragment ions agree with previously reported masses for ornicorrugatin (Matthijs, et al., 2008).

| Fragment composition | Experimental mass (*m/z*) | Calculated mass (m/z) |
|---|---|---|
| $C_{19}H_{34}N_5O_3^+$ | 380.2681 | 380.2662 |
| $C_{23}H_{38}N_7O_4^+$ | 476.3011 | 476.2985 |
| $C_{31}H_{57}N_{11}O_8^+$ | 711.4194 | 711.4392 |
| $C_{35}H_{60}N_{11}O_{13}^+$ | 842.4422 | 842.4372 |
| $C_{37}H_{62}N_{11}O_{16}^+$ | 916.4429 | 916.4376 |
| $C_{41}H_{62}N_{13}O_{15}^+$ | 976.4544 | 976.4488 |
| Ornicorrugatin M+H, $C_{41}H_{66}N_{13}O_{17}^+$ | 1012.4760 | 1012.4699 |

**Figure S15. UPLC-MS spectrum of pyoverdine siderophore from *Pseudomonas brassicacearum* DSM 13227.** The singly charged mass of *m/z* 1134.4 Da $[M+H]^+$ and the doubly charged mass of *m/z* 567.7 Da $[M+2H]^{2+}$ are consistent with the *m/z* values of the pyoverdine, m/z 1134.4339 Da $[M+H]^+$.

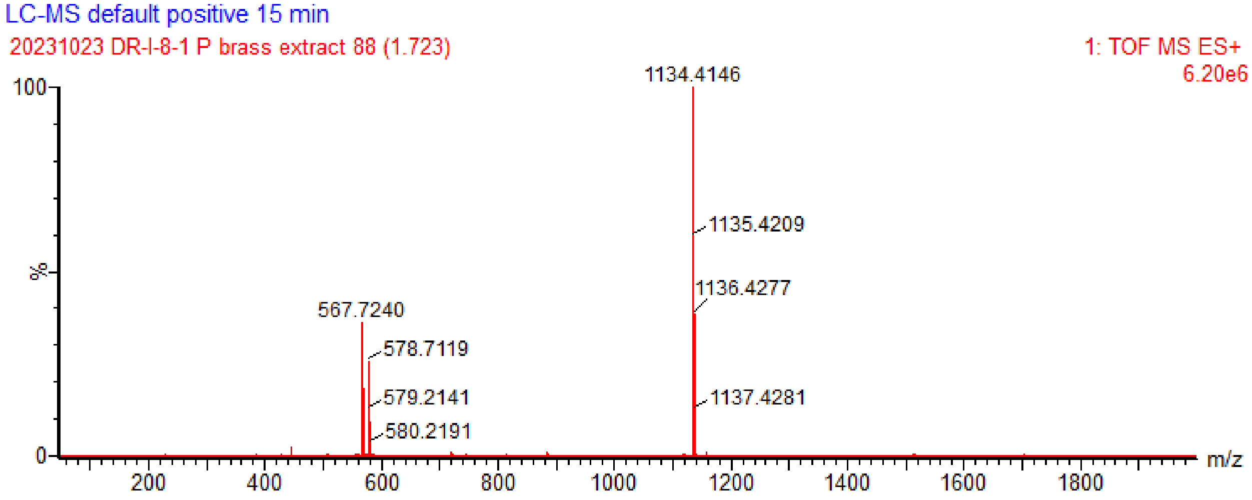

**Figure S16. LC-MS tandem MS spectrum of pyoverdine siderophore, pyoverdine A214, from *Pseudomonas brassicacearum* DSM 13227.**

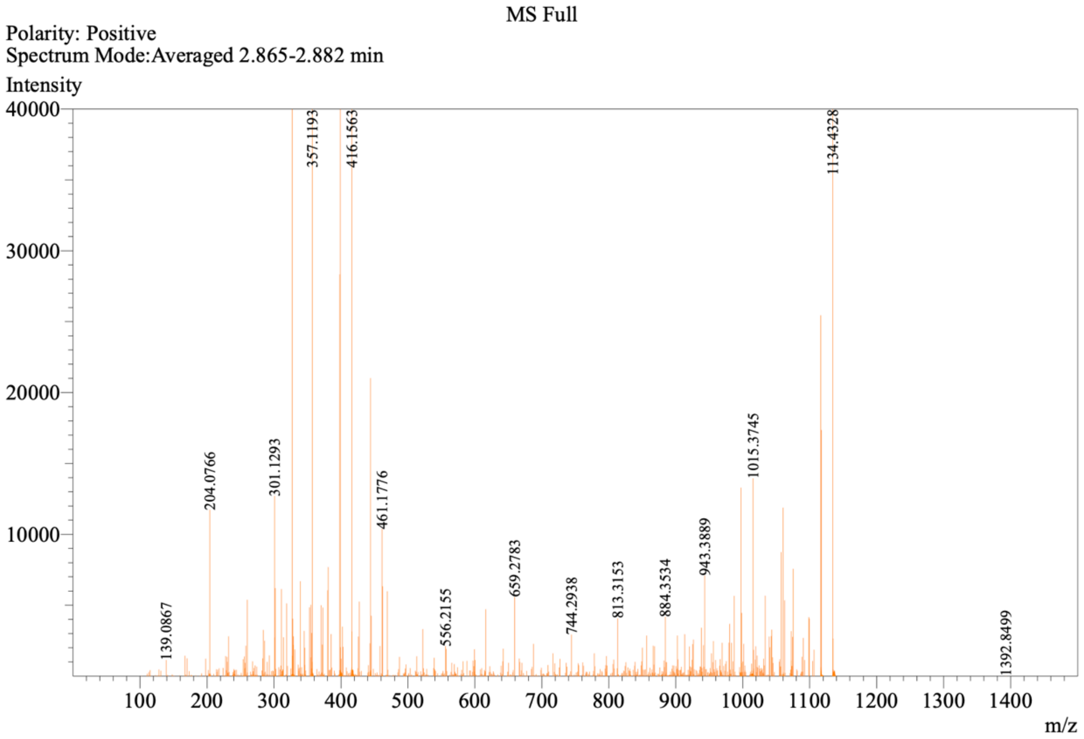
